# Vapor inhalation of cannabidiol (CBD) in rats

**DOI:** 10.1101/659250

**Authors:** Mehrak Javadi-Paydar, Kevin M. Creehan, Tony M. Kerr, Michael A. Taffe

**Author notes:** Address Correspondence to: Dr. Michael A. Taffe, Department of Psychiatry, Mail Code 0714; 9500 Gilman Drive; University of California, San Diego, La Jolla, CA 92093; USA.

## Abstract

Cannabidiol (CBD) is increasingly available in e-cigarette liquids and other products. CBD use has been promoted for numerous purported benefits which have not been rigorously assessed in preclinical studies. The objective of this study was to further validate an inhalation model to assess CBD effects in the rat. The primary goal was to determine plasma CBD levels after vapor inhalation and compare that with the levels observed after injection. Secondary goals were to determine if hypothermia is produced in male Sprague-Dawley rats and if CBD affects nociception measured by the warm water tail-withdrawal assay. Blood samples were collected from rats exposed for 30 minutes to vapor generated by an e-cigarette device using CBD (100, 400 mg/mL in the propylene glycol vehicle). Separate experiments assessed the body temperature response to CBD in combination with nicotine (30 mg/mL) and the anti-nociceptive response to CBD. Vapor inhalation of CBD produced concentration-related plasma CBD levels in male and female Wistar rats that were within the range of levels produced by 10 or 30 mg/kg, CBD, i.p.. Dose-related hypothermia was produced by CBD in male Sprague-Dawley rats and this was partially attenuated by 5-HT1a receptor blockade. Nicotine (30 mg/mL) inhalation enhanced the effect of CBD. CBD inhalation had no effect on anti-nociception alone or in combination with Δ^9^-tetrahydrocannabinol inhalation. The vapor-inhalation approach is a suitable pre-clinical model for the investigation of the effects of inhaled CBD. This route of administration produces hypothermia in rats, while i.p. injection does not at comparable plasma CBD levels.

## Introduction

Cannabidiol (CBD), a cannabinoid compound found in many strains of the *Cannabis* genus, has experienced recent popularity for claimed beneficial effects on health and well-being (Lewis, 2019). The past few years has witnessed increasing availability of e-cigarette liquids represented as containing only CBD (Peace et al., 2016) and indeed many do contain only CBD with only a subset appearing to contain other psychoactive drugs (Poklis et al., 2019). Another popular set of CBD products purport to deliver the drug transdermally via oils and creams and yet another set of products deliver CBD orally via gummy candies and other edible items. The Agriculture Improvement Act of 2018 which was signed into law in the USA in December of 2018 expanded legal hemp cultivation for the production of cannabidiol (Bourque, 2018). CBD use has been promoted by social and traditional media “lifestyle” celebrities, including for anxiety related to impending child birth (Haller, 2019). Thus, the use of CBD is likely to continue to increase.

These recent and growing CBD use trends prompt renewed interest in preclinical study of the potential impact of CBD, both to test claims of beneficial effects on health and well-being and to identify any likely risks. Pre-clinical experiments typically inject rodents with CBD i.p. or s.c. which may not reflect the pharmacokinetic profile observed after vapor inhalation. Despite growing popularity there is a deficit of pre-clinical information on the breadth of potential benefits and harms, as recently reviewed (Boggs et al., 2018). Although sometimes termed non-psychoactive due to a lack of many of the effects associated with Δ^9^-tetrahydrocannabinol, it has been shown more recently to have pharmacological effects in humans and in pre-clinical models. CBD has been approved for treatment of two seizure disorders in the US (Sekar and Pack, 2019) and has been approved, in combination with THC, for spasticity associated with multiple sclerosis in Canada (Keating, 2017; Oreja-Guevara, 2012). A recent paper has shown transdermal efficacy in rat models of cocaine and alcohol addiction (Gonzalez-Cuevas et al., 2018), this route of administration aligns more closely with another very common route of CBD use at present via creams and oils.

CBD produces at least some of its effects, *in vivo*, by acting as an agonist at the serotonin 1A (5-HT_1A_) receptor subtype. For example, intravenous CBD dose-dependently decreases the firing rate of 5-HT neurons in dorsal raphe region in a manner that can be prevented by the 5-HT_1A_ antagonist WAY 100,635 but not the CB1 antagonist AM251 (De Gregorio et al., 2019). Anti-emetic and anti-nausea-like effects of CBD are likewise attenuated by WAY 100,635 (Rock et al., 2012) and CBD blocks hyperphagia caused by the 5-HT_1A_ agonist 8-OH-DPAT (Scopinho et al., 2011). The 5-HT_1A_ agonist 8-OH-DPAT produces profound hypothermia in rats, while the 5-HT_1A_ antagonist WAY 100,635 attenuates these effects, thus it is peculiar that cannabidiol does not appear to reduce body temperature when injected in rats (Bloom et al., 1978; Hiltunen et al., 1988; Taffe et al., 2015) or mice (Hayakawa et al., 2008). We recently showed that vapor inhalation of CBD reduced the body temperature of male and female rats (Javadi-Paydar et al., 2018a), which can be directly contrasted with a failure to observe any temperature change in rats following i.p. injection of CBD at 30-60 mg/kg doses using similar methods to evaluate body temperature responses (Taffe et al., 2015). The inhalation study used an e-cigarette based vapor inhalation approach which has now been shown effective for delivering physiologically relevant doses of sufentanil, THC, methamphetamine, the substituted cathinones (“bath salts”) MDPV and 4-MMC as well as nicotine to rats (Javadi-Paydar et al., 2019; Nguyen et al., 2016a; Nguyen et al., 2016b; Vendruscolo et al., 2018).

Gonzalez-Cuevas and colleagues (Gonzalez-Cuevas et al., 2018) reported plasma CBD levels 4 hours after application of 15 and 30 mg/kg doses using a dermal CBD gel. It is possible that the i.p. injection of 30-60 mg/kg CBD does not result in plasma exposure similar to that achieved with vapor inhalation (Javadi-Paydar et al., 2018a) or with a dermal gel application, potentially explaining a differential efficacy. The present study was therefore conducted primarily to determine if the plasma CBD levels obtained following vapor inhalation and i.p. injection differed substantially. Since our prior injection study (Taffe et al., 2015) was in Sprague-Dawley rats and the inhalation study (Javadi-Paydar et al., 2018a) was in Wistar rats, this study further set out to determine if vapor inhalation of CBD reduces the body temperature of Sprague-Dawley males. We previously observed a greater sensitivity of Sprague-Dawley male rats compared with Wistar male rats to hypothermia induced by *THC* inhalation despite similar plasma THC levels and similar antinociceptive responses in a direct comparison study (available in pre-print form: https://www.biorxiv.org/content/10.1101/541003v1), thus it appeared critical to determine whether any strain differences in response to CBD. Mechanistic studies were included to determine if WAY 100,635 attenuates the hypothermia generated by CBD inhalation.

Finally, this investigation included a study of combined CBD/nicotine inhalation to compare with our recent study (Javadi-Paydar et al., 2019) demonstrating the effects of combined THC and nicotine inhalation on body temperature (additive) and locomotor activity (subtractive). Tucker and colleagues report that 37% of a young adult sample co-use tobacco / nicotine with marijuana (Tucker et al., 2019). It has recently been shown that CBD may reduce methamphetamine seeking (Hay et al, 2018) and alcohol and cocaine seeking (Gonzalez-Cuevas, et al, 2018) in rats. CBD may also reduce bias to tobacco-related cues during smoking abstinence in humans (Hindocha et al., 2018b) and a preliminary double-blind placebo controlled study found that CBD delivered by inhaler was able to reduce cigarette consumption over a week of treatment (Morgan et al., 2013). CBD also attenuates memory impairment associated with nicotine withdrawal in mice (Saravia et al 2019) but does not alleviate cognitive impairment associated with nicotine abstinence in human smokers (Hindocha et al., 2018a).

## 2. Methods

### 2.1 Subjects

Female and male Wistar rats (N=8 per group; Charles River) and male Sprague-Dawley (N=7; Harlan/Envigo, Livermore, CA) rats were housed in humidity and temperature-controlled (23±2 °C) vivaria on 12:12 hour light:dark cycles. Rats had *ad libitum* access to food and water in their home cages and all experiments were performed in the rats’ scotophase. Wistar rats entered the laboratory on PND 22 and the Sprague-Dawley rats entered the laboratory at 10-11 weeks of age. All procedures were conducted under protocols approved by the Institutional Animal Care and Use Committee of The Scripps Research Institute.

### 2.2 Inhalation Apparatus

Sealed exposure chambers were modified from the 259mm × 234mm × 209mm Allentown, Inc (Allentown, NJ) rat cage to regulate airflow and the delivery of vaporized drug to rats using e-cigarette cartridges (Protank 3 Atomizer, MT32 coil operating at 2.2 ohms, by Kanger Tech; Shenzhen Kanger Technology Co.,LTD; Fuyong Town, Shenzhen, China; controlled by e-vape controller Model SSV-1; La Jolla Alcohol Research, Inc, La Jolla, CA, USA) or (SMOK® TFV8 with X-baby M2 atomizer; 0.25 ohms dual coil; Shenzhen IVPS Technology Co., LTD; Shenzhen, China; controlled by e-vape controller Model SSV-3; La Jolla Alcohol Research, Inc, La Jolla, CA, USA) as has been previously described (Javadi-Paydar et al., 2019; Javadi-Paydar et al., 2018a; Nguyen et al., 2016b). The controllers were triggered to deliver the scheduled series of puffs by a computerized controller designed by the equipment manufacturer (Control Cube 1; La Jolla Alcohol Research, Inc, La Jolla, CA, USA). The chamber air was vacuum controlled by a chamber exhaust valve (i.e., a “pull” system) to flow room ambient air through an intake valve at ~1 L per minute. This also functioned to ensure that vapor entered the chamber on each device triggering event. The vapor stream was integrated with the ambient air stream once triggered.

### 2.3 Drugs

Rats were exposed to vapor generated from cannabidiol (CBD; 100, 200, 400 mg/mL), Δ^9^-tetrahydrocannabinol (THC; 25, 50, 100 mg/mL) or nicotine bitartrate (30 mg/mL) dissolved in a propylene glycol (PG) vehicle. Inhalation concentrations for CBD and nicotine were derived from our prior studies (Javadi-Paydar et al., 2019; Javadi-Paydar et al., 2018a). For the Protank 3 / SSV-1 apparatus, four 10-s vapor puffs were delivered with 2-s intervals every 5 minutes. For the SMOK / SSV-3 apparatus, one 6-s vapor puff was delivered every 5 minutes. CBD (10, 30 mg/kg) was injected, i.p., in a 1:1:8 ratio of ethanol:cremulphor:saline and WAY 100,635 was injected, i.p., in saline. Nicotine, 8-hydroxy-2-(dI-n-propylamino)tetralin (8-OH-DPAT), SB 269970 and WAY 100,635 were obtained from Sigma-Aldrich (St. Louis, MO). The WAY 100,635 doses (0.1, 1.0 mg/kg, i.p.) were selected from prior studies demonstrating efficacy against thermoregulatory disruption produced by 8-OH-DPAT in rats (Mensonides-Harsema et al., 2000; Nicholas and Seiden, 2003).

### 2.4 Radiotelemetry

Sprague-Dawley rats (N=15) were anesthetized with an isoflurane/oxygen vapor mixture (isoflurane 5% induction, 1-3% maintenance) and sterile radiotelemetry transmitters (Data Sciences International, St. Paul, MN; TA-F40) were implanted in the abdominal cavity through an incision along the abdominal midline posterior to the xyphoid space as previously described (Javadi-Paydar et al., 2018b). Activity and temperature responses were evaluated in separate recording chambers (Experiment 2) or in the vapor inhalation chambers (Experiment 3) in a dark testing room, separate from the vivarium, during the (vivarium) dark cycle. Data were collected on a 5-minute schedule throughout the recording intervals.

### 2.5 Experiments

#### 2.5.1 Experiment 1: CBD Plasma Concentrations After Vapor Inhalation and Injection

Groups of female (N=8; 338.6 g, SEM 9.7, bodyweight; 38 weeks of age) and male (N=8; 653.1g, SEM 21.1, bodyweight; 38 weeks of age) Wistar rats were used in this study. This group had been exposed to THC vapor on PND 31, for 30 minutes twice daily from PND 36-39 (Nguyen et al., 2018), and on PND 86, PND 100, PND 107. The animals were also exposed to nicotine (30 mg/mL) vapor inhalation and nicotine injection on PND 212 and PND 226. All of the foregoing studies involved obtaining blood samples once or twice after drug administration. For the current study, blood (~300 µl) was withdrawn at 35 and 120 minutes after the start of 30-minute CBD (100, 400 mg/mL in the PG vehicle) sessions by acute venipuncture of the jugular vein, using a 25 gauge needle under isoflurane inhalation anesthesia. Rats were permitted to recover after the first sample and then re-anesthetized for the second sample. Samples were also withdrawn at 35 and 120 minutes after the injection of CBD (10, 30 mg/kg, i.p.). A minimum of two weeks recovery was interposed between each experiment.

#### 2.5.2 Experiment 2: CBD Effects on Body Temperature and Activity

A group (N=8; 664 g, 44.7 SEM, bodyweight; 33 weeks of age) of male Wistar rats implanted with radiotelemetry devices and used previously to determine effects of MDMA by vapor inhalation and i.p. injection (5-6 total active drug sessions, no more than twice per week) were used in studies to confirm that CBD decreases temperature at these inhalation concentrations. Animals were exposed to inhalation of PG or CBD (100 mg/mL; 30 minutes) in a counter balanced order and then three weeks later to inhalation of PG or CBD (400 mg/mL) in a counter balanced order. A followup study was conducted to determine if the thermoregulatory effects of CBD could be attenuated by the 5-HT_1A_ antagonist WAY 100,635. For this study rats were exposed to 30 minutes of inhalation of CBD (100 mg/mL) or THC (100 mg/mL) with 0.0, 0.1 or 1.0 mg/kg WAY 100,635 administered 15 minutes before vapor initiation, in a counter balanced order. Finally, a positive control experiment was conducted to determine if injection of the 5-HT_1A_ agonist 8-OH-DPAT (0.1 mg/kg, i.p.), would decrease body temperature and if such effects were attenuated by pre-treatment with the 5-HT_1A_ antagonist WAY 100,635 or the 5-HT_7_ antagonist SB-269970 (Faure et al., 2006; Hedlund et al., 2004; Wright et al., 2012).

#### 2.5.3 Experiment 3: CBD and Nicotine Effects on Body Temperature and Activity

A group (N=7; 551.4 g, SEM 9.0, bodyweight; 49 weeks of age) of male Sprague-Dawley rats implanted with radiotelemetry devices and used previously to determine effects of nicotine vapor inhalation alone and in combination with THC (Javadi-Paydar et al., 2019) were used in studies to confirm that CBD decreases temperature in this strain. Animals were first exposed to inhalation of PG, Nicotine (30 mg/mL), CBD (400 mg/mL) or the combination for 30 minutes in a counter balanced order. For reference, the inhalation of Nicotine (30 mg/mL) for 30 minutes was shown in the prior study (Javadi-Paydar et al., 2019) to produce post-session mean plasma nicotine of 61 ng/mL (SEM 7.8) and mean plasma cotinine of 32.4 ng/mL (SEM 6.2). Then, to determine if the effects of CBD were dose dependent, the 4 conditions were repeated with only the CBD inhalation concentration changed (100 mg/mL). Animals were next exposed to inhalation of CBD (100 mg/mL) with injection of the antagonist WAY 100635 (0.0, 0.1, 1.0 mg/kg, i.p.) 15 minutes prior to starting inhalation, in a counter balanced order. A minimum 3-4 day interval was interposed between active dosing experiments.

**Figure 1:**
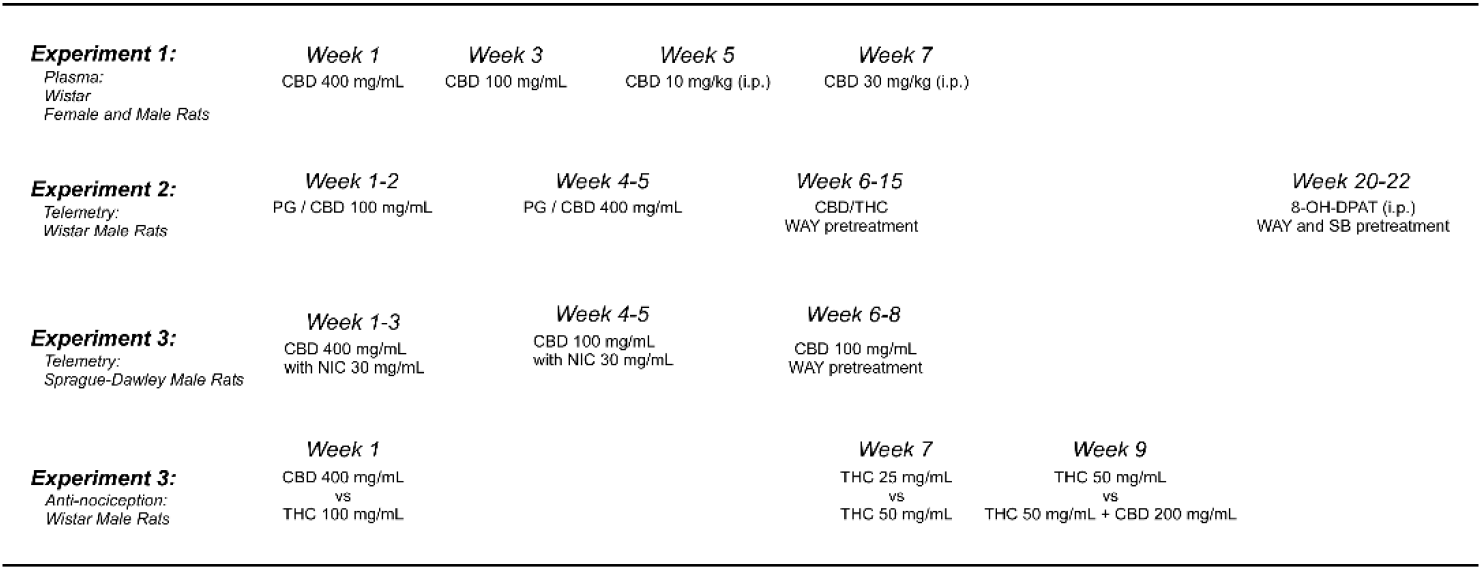
Timeline depicting the course of experiments conducted in each cohort of rats.

#### 2.5.4 Experiment 4: CBD Effects on THC-induced Anti-nociception

A group (N=6; 728.6 g, SEM 35.3, bodyweight; 47 weeks of age) of male Wistar rats were used to determine if vapor inhalation of CBD would modify the anti-nociceptive effects of THC using a warm-water tail-withdrawal assay shown previously to be sensitive to THC vapor inhalation in male and female rats (Javadi-Paydar et al., 2018a; Nguyen et al., 2016b). Three separate two-dose condition studies were conducted with tail-withdrawal evaluated before the inhalation session and then 5 min after the 30-minute session. For each study the two inhalation conditions were evaluated in a counterbalanced order and a 3-4 day interval was interposed between inhalation session. The first experiment contrasted the effect of CBD (400 mg/mL) and THC (100 mg/mL). Then to determine a minimal effective dose, the next experiment contrasted the effects of THC 25 vs 50 mg/mL. Finally, the effect of CBD (200 mg/mL) + THC (50 mg/mL) versus THC (50 mg/mL) alone was determined.

### 2.7 Data Analysis

Plasma levels of CBD were analyzed with ANOVA with repeated measures factors of Drug Condition and Time post-initiation of vapor or Time post-injection. A between-subjects factor was included for sex. Temperature and activity rate (counts per minute) data were collected via the radiotelemetry system on a 5-minute schedule and analyzed in 30 minute averages (the time point refers to the ending time, i.e. 60 = average of 35-60 minute samples). In the WAY pre-treatment study, the baseline interval is depicted as two 15-minute bins with the i.p. injection just prior to the second baseline interval. Any missing temperature values were interpolated from surrounding values, this amounted to less than 10% of data points. Telemetry data were analyzed with Analysis of Variance (ANOVA) with repeated measures factors for the Drug Condition and the Time post-initiation of vapor or Time post-injection. Comparison with the pre-treatment baselines and the vehicle conditions. The tail withdrawal latencies were analyzed with within-subjects factors for pre/post-inhalation and for inhalation condition. Any significant effects within group were followed with post-hoc analysis using Tukey correction for all multi-level, and Sidak correction for any two-level, comparisons. All analysis used Prism 7 or 8 for Windows (v. 7.03, 8.1.1; GraphPad Software, Inc, San Diego CA).

## 3. Results

### 3.1 Experiment 1: CBD Plasma Concentrations After Vapor Inhalation and Injection

Vapor inhalation produced CBD concentration dependent effects on plasma CBD (Figure 2) in male and female Wistar rats. Three-way analysis of the vapor inhalation data confirmed a significant difference in plasma CBD attributable to Sex [F (1, 14) = 7.34; P<0.05], to the CBD concentration [F (1, 14) = 35.82; P<0.0001], to the time of sampling [F (1, 14) = 171.6; P<0.0001] and the interaction of sex with Time [F (1, 14) = 4.62; P<0.05]. Collapsed across sex, the post-hoc test confirmed significant differences across all six possible comparisons.

**Figure 2:**
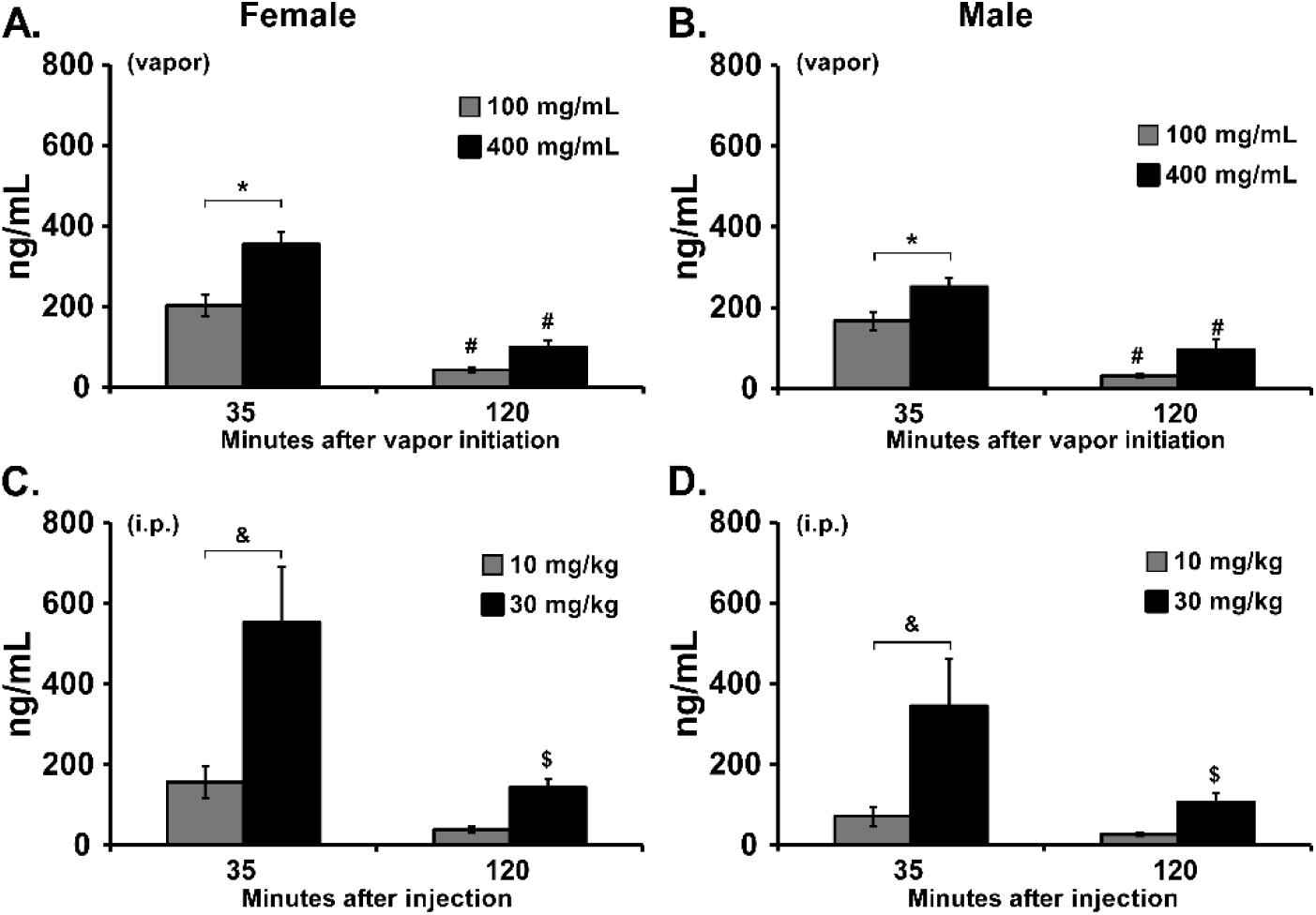
A, B) Mean plasma concentration of CBD following vapor inhalation for 30 minutes in groups of A) female (N=8), and B) male (N=8), rats. A significant difference between concentrations, within each sex, is indicated with *, and a significant difference between timepoints after vapor initiation is indicated with #. C, D) Mean plasma concentration of CBD following i.p. injection in the same groups. Collapsed across sex, a significant difference between doses is indicated with &, and a significant difference between timepoints after vapor initiation with $.

To further explore the effects within sex groups, two-way ANOVAs were conducted and confirmed significant differences in plasma CBD for female [Vape Concentration, F (1, 7) = 10.86; P<0.05; Time, F (1, 7) = 165.0; P<0.0001; Interaction, F (1, 7) = 5.46; P=0.052] and male [Vape Concentration, F (1, 7) = 14.72; P<0.01; Time, F (1, 7) = 310.8; P<0.0001; Interaction, F (1, 7) = 0.35; P=0.5712] rats. The post-hoc tests confirmed a significant concentration related difference in plasma CBD immediately after the inhalation session and a significant reduction 120 minutes after the start of inhalation. Differences in plasma CBD were also observed in the intraperitoneal injection study. The three-way ANOVA confirmed significant effects of Time after injection [F (1, 14) = 19.76; P<0.001], CBD Dose [F (1, 14) = 20.96; P<0.001] and the interaction of Dose with Time [F (1, 14) = 6.74; P<0.05]. Collapsed across sex, the post-hoc test confirmed significant differences between doses at 35 minutes after injection and a difference between time points for the 30 mg/kg, i.p., dose.

### 3.2 Experiment 2: CBD Effects on Body Temperature and Activity

CBD produced a significant temperature reduction in male Wistar rats following inhalation at the 100 or 400 mg/mL concentrations (Figure 3). The ANOVA confirmed significant effects of Time after vapor initiation [F (7, 49) = 12.92; P<0.0001], of Vape Condition [F (3, 21) = 16.49; P<0.0001] and the interaction of factors [F (21, 147) = 15.01; P<0.0001] on body temperature. The post-hoc test further confirmed that temperature was lower than the baseline and the respective vehicle inhalation condition 60-120 minutes after CBD (100 mg/mL) inhalation and 60-180 minutes after CBD (400 mg/mL) inhalation. The analysis of activity rates confirmed significant The ANOVA confirmed significant effects of Time after vapor initiation [F (7, 49) = 23.59; P<0.0001], of Vape Condition [F (3, 21) = 5.88; P<0.005] and the interaction of factors [F (21, 147) = 1.925; P<0.05] on body temperature. The post-hoc test confirmed that activity rate was significantly lower, compared with the other inhalation conditions, 60 minutes after the initiation of CBD (400 mg/mL) inhalation.

**Figure 3:**
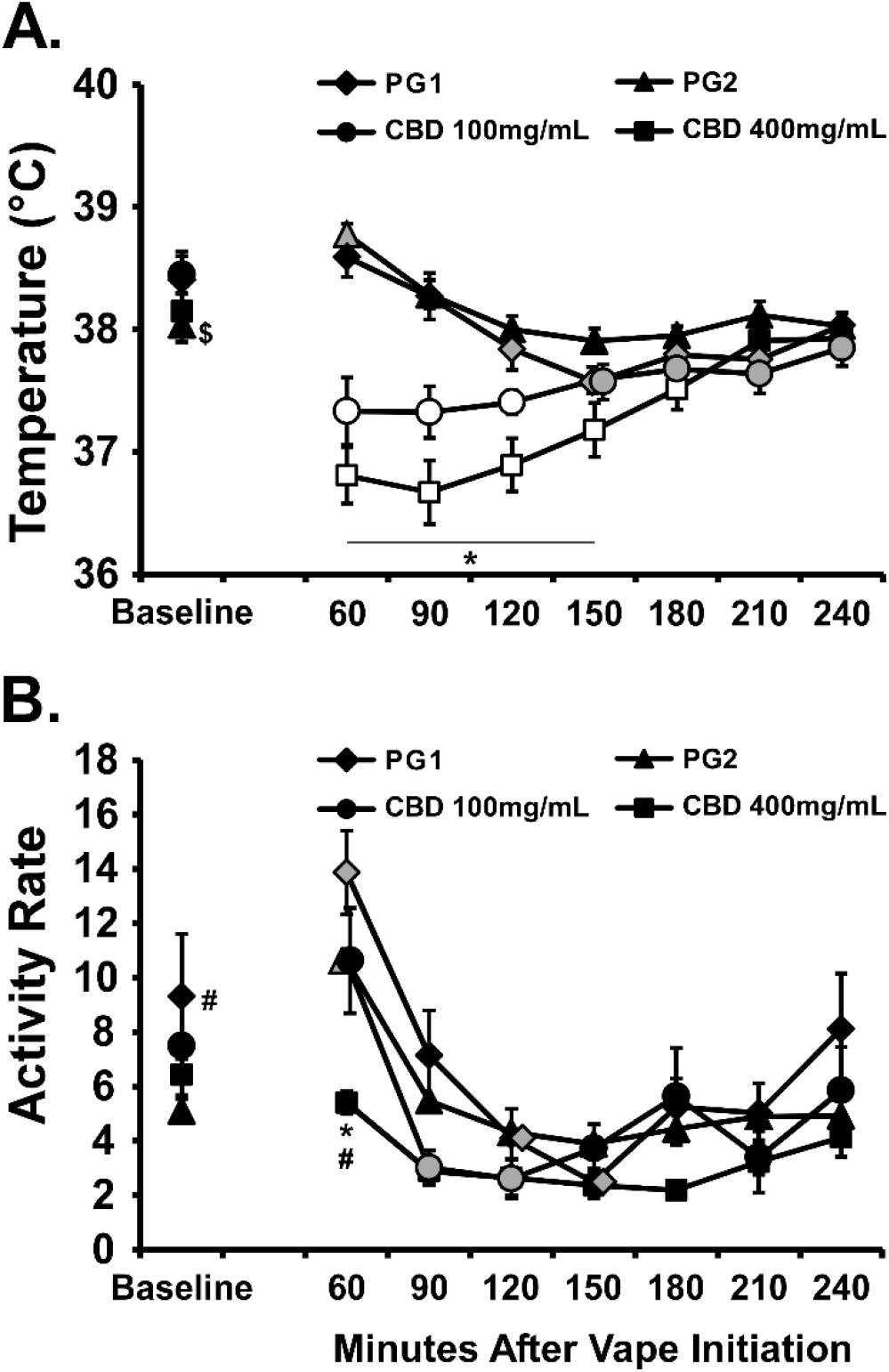
Mean (N=8; SEM) body temperature and activity following inhalation of the PG vehicle, cannabidiol (CBD; 100, 400 mg/mL) for 30 minutes. Shaded symbols indicate a significant difference from the Baseline (Base), within treatment condition and open symbols represent a significant difference from the baseline and the respective time-point after PG inhalation. A significant difference between CBD (100 mg/mL) and CBD (400 mg/mL) indicated by *, a significant difference from PG2 with # and a difference from PG1 with $.

The 5-HT_1A_ antagonist WAY 100,635 did not significantly alter the thermoregulatory response to the inhalation of CBD or THC (Figure 4). The ANOVA confirmed that there were significant effects of dosing condition [F (5, 35) = 7.54; P<0.0001], of time after vapor initiation [F (7, 49) = 9.02; P<0.0001] and the interaction [F (35, 245) = 1.55; P<0.05] on body temperature. The post-hoc test confirmed that temperature was reduced below baseline in all of the THC conditions, and in the Sal-CBD and 0.1 WAY-CBD conditions. Nevertheless, there were no differences between the saline pre-treatment and the WAY pre-treatment conditions for either THC or CBD inhalation. The ANOVA confirmed that there were significant effects of dosing condition [F (5, 35) = 3.98; P<0.01] and of time after vapor initiation [FF (7, 49) = 40.86; P<0.0001 on activity rate.

**Figure 4:**
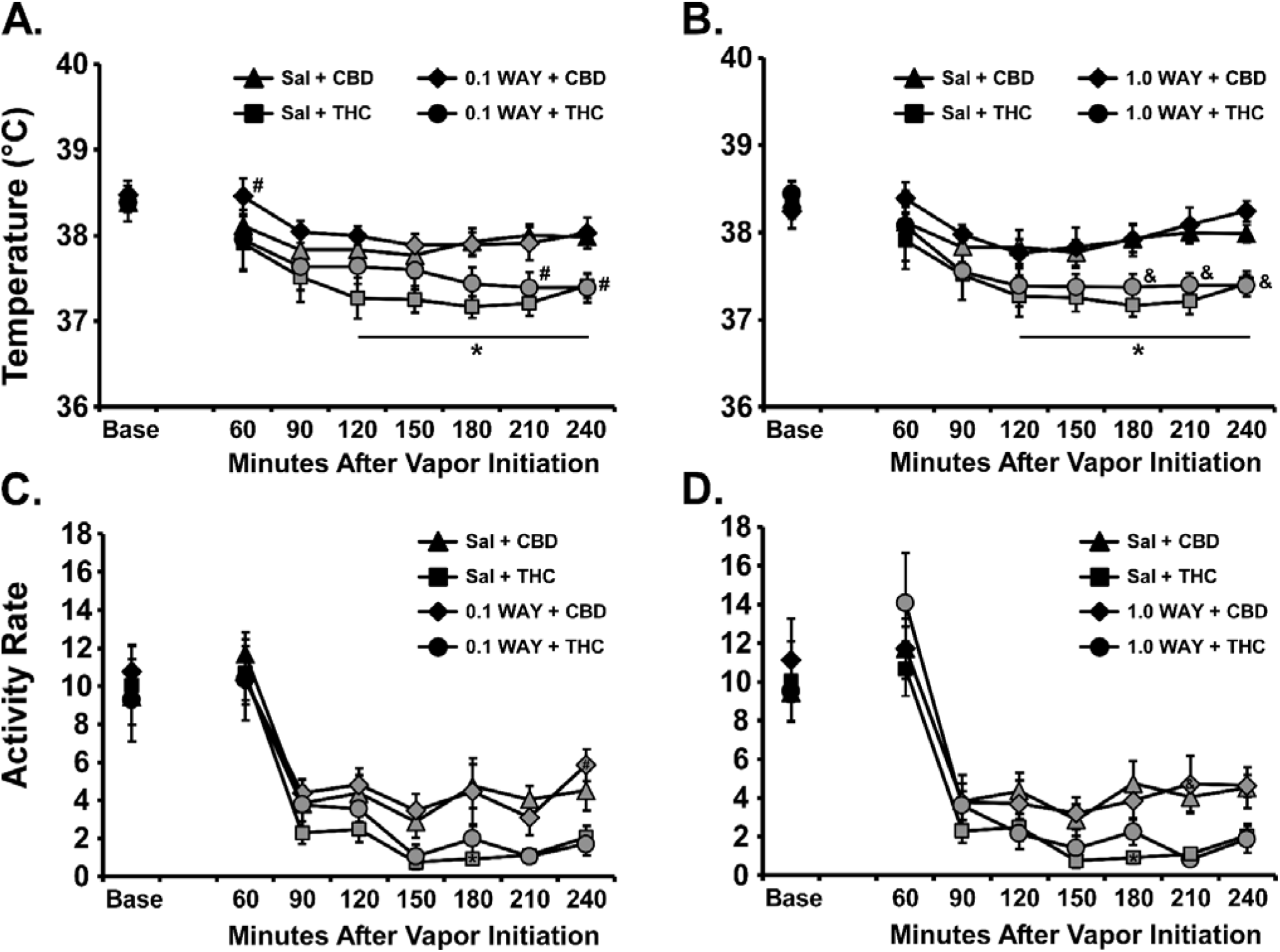
Mean (N=8; SEM) A, B) body temperature and C, D) activity following inhalation of cannabidiol CBD (100 mg/mL) or THC (100 mg/mL) with pre-inhalation injection with WAY 100,635 (0.1, 1.0 mg/mg, i.p.). Shaded symbols indicate a significant difference from the Baseline (Base), within treatment condition A significant difference between CBD and THC with saline pretreatment is indicated by *, with 0.1 mg/kg WAY pretreatment by # and with 1.0 mg/kg WAY pretreatment by &.

The positive control experiment in this group found that the 5-HT_1A_ agonist 8-OH-DPAT produced hypothermia when injected, and that the 5-HT_1A_ antagonist WAY 100,635 (0.1, 1.0 mg/kg, i.p.) or 5-HT_7_ antagonist SB-269970 (0.1, 1.0 mg/kg, i.p.) can attenuate the response (Figure 5). In the <u>WAY 100,635 experiment</u> the ANOVA confirmed that there were significant effects of Dosing Condition [F (2, 14) = 30.99; P<0.0001], of Time after 8-OH-DPAT injection [F (8, 56) = 32.56 P<0.0001] and the interaction of factors [F(16, 112) = 5.28; P<0.0001] on body temperature. The post-hoc test confirmed significant differences between the Saline pre-treatment condition and each of the 0.1 mg/kg (30-180 minutes after 8-OH-DPAT injection) and 0.1 mg/kg (30-150 minutes after 8-OH-DPAT injection) doses of WAY 100,635. There was a significant effect of Time post-injection [F (8, 56) = 33.18; P<0.0001], but not of Dosing Condition or of the interaction of factors on activity rate. In the <u>SB-269970 experiment</u> the ANOVA confirmed that there were significant effects of Dosing Condition [F (2, 14) = 19.03; P=0.0001], of Time after 8-OH-DPAT injection[F (8, 56) = 21.69; P<0.0001] and the interaction of factors [F (16, 112) = 3.16; P<0.0005] on body temperature.

**Figure 5:**
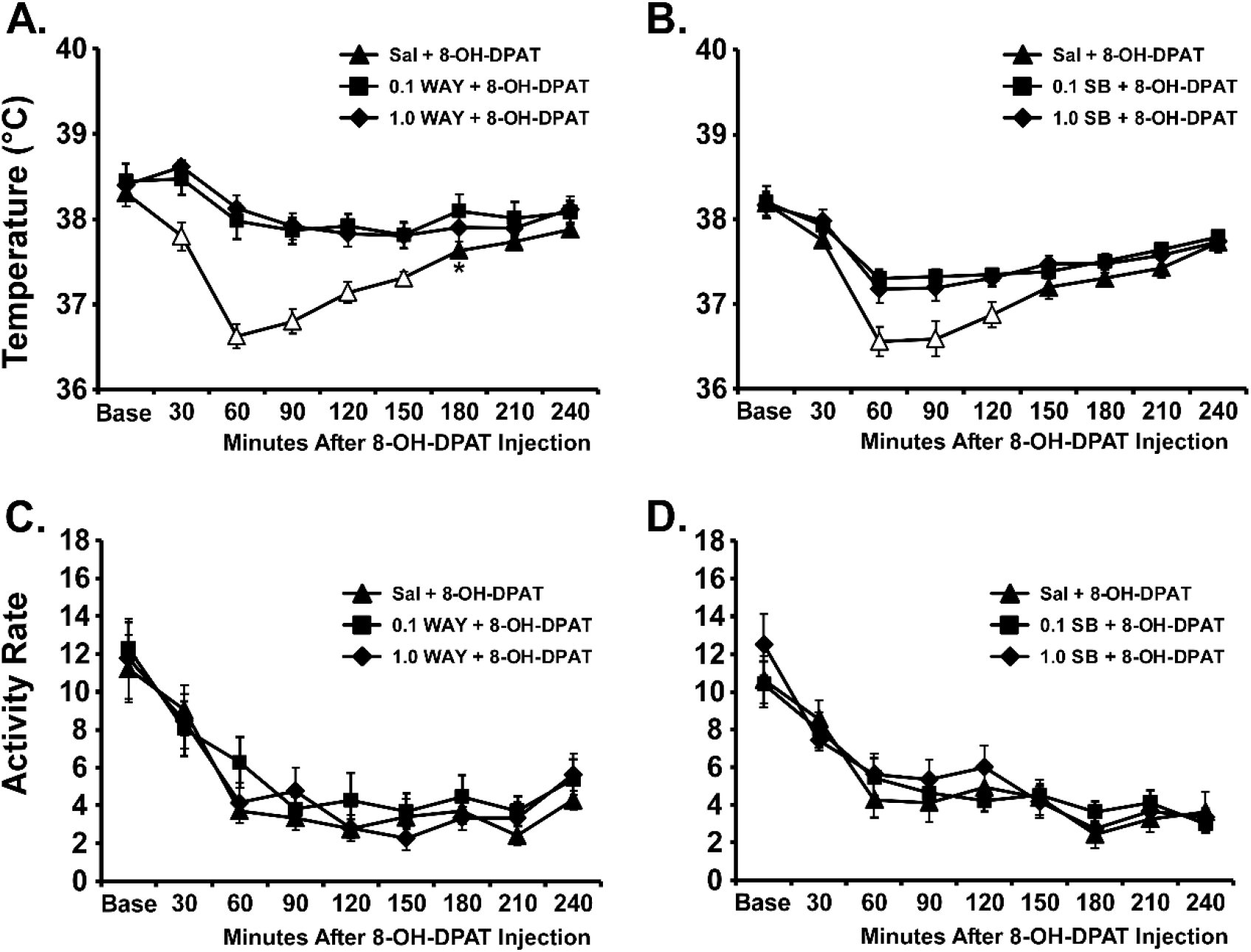
Mean (N=8; SEM) A, B) body temperature and C, D) activity rates following injection of 8-OH-DPAT (0.1 mg/kg, i.p.) 15 minutes after pre-treatment with A, C) WAY-100,635 (0.0, 0.1, 1.0 mg/mg, i.p.) or B,D) SB-269970 (0.0, 0.1, 1.0 mg/mg, i.p.). Open symbols indicate a significant difference from the Baseline (Base) within treatment condition and from each of the other pre-treatments at the respective time post-injection. A significant difference between Saline and 0.1 mg/kg WAY pretreatment is indicated with *.

The post-hoc test confirmed significant differences between the Saline pre-treatment condition and each of the 0.1 mg/kg (60-120 minutes after 8-OH-DPAT injection) and 0.1 mg/kg (60-120 minutes after 8-OH-DPAT injection) doses of SB-269970. There was also a significant effect of Time post-injection [F (8, 56) = 34.54; P<0.0001] on activity rate.

### 3.3 Experiment 3: CBD and Nicotine Effects on Body Temperature and Activity

#### 3.3.1. CBD and Nicotine Alone and In Combination

CBD again produced a significant temperature reduction following inhalation at the 100 or 400 mg/mL concentrations (Figure 6A,B), in this case in male Sprague-Dawley rats. Nicotine (30 mg/mL) had no effect by itself but enhanced the hypothermia induced by either concentration of CBD. The ANOVA confirmed significant effects of Time after vapor initiation (F (7, 42) = 131.8; P<0.0001), of Vape Condition (F (7, 42) = 19.39; P<0.0001) and the interaction of factors (F (49, 294) = 7.52; P<0.0001). No differences in body temperature were confirmed for any timepoint after inhalation of the PG or Nicotine (30 mg/mL) conditions for either dose series. The post-hoc test did confirm a significant difference between the temperatures observed after the CBD (100 mg/mL) and CBD (400 mg/mL) inhalation conditions from 60-210 minutes after vapor initiation, and between the CBD (100 mg/mL) + Nicotine (30 mg/mL) and CBD (400 mg/mL) + Nicotine (30 mg/mL) conditions from 120-210 minutes after vapor initiation. Furthermore, body temperature differed significantly between the CBD (100 mg/mL) and CBD (100 mg/mL) + Nicotine (30 mg/mL) conditions from 60-150 minutes after vapor initiation and between the CBD (400 mg/mL) and CBD (400 mg/mL) + Nicotine (30 mg/mL) conditions at 120 minutes after vapor initiation.

**Figure 6:**
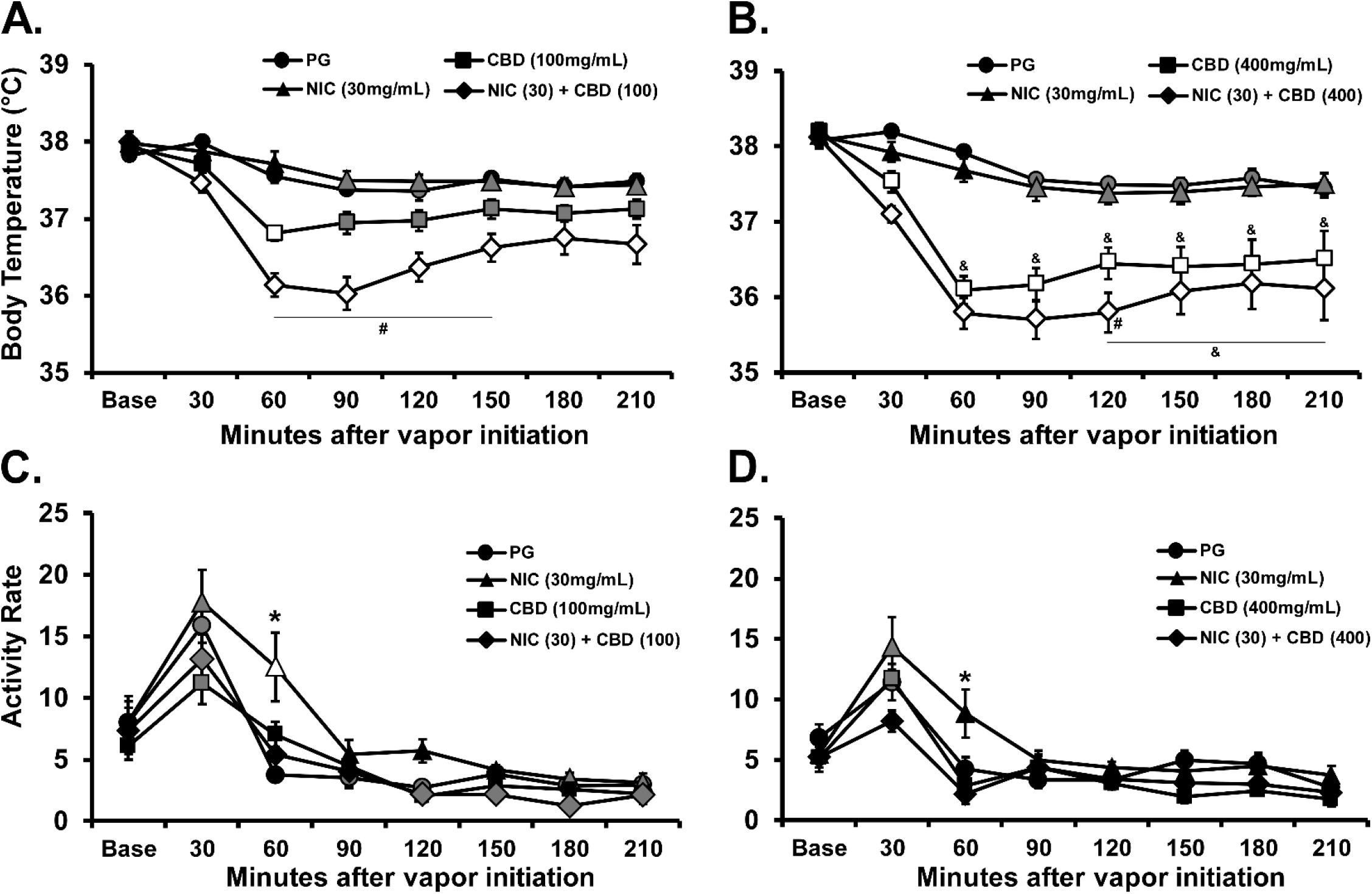
Mean (N=7; SEM) A, B) body temperature and C, D) activity following inhalation of the PG vehicle, cannabidiol (CBD), Nicotine (NIC; 30 mg/mL) or the CBD/NIC combination for 30 minutes. The panels present the experiments with the A, C) CBD (100 mg/mL) and B, D) CBD (400 mg/mL) concentrations. Shaded symbols indicate a significant difference from the Baseline (Base), within treatment condition and open symbols represent a significant difference from the baseline and the respective time-point after PG inhalation. A significant difference from all other treatment conditions at a given time-point is indicated with *, a significant difference across CBD concentration is indicated with & and a significant difference between CBD alone and CBD+ Nicotine is indicated with #.

Analysis of the locomotor activity (Figure 6C,D) confirmed significant effects of Time after vapor initiation (F (7, 42) = 52.45; P<0.0001), of Vape Condition (F (7, 42) = 6.22; P<0.0001) and the interaction of factors (F (49, 294) = 2.36; P<0.0001). The post-hoc test confirmed a significant difference between activity observed 60 minutes after vapor initiation in the Nicotine (30 mg/mL) versus all other conditions in each dose series.

#### 3.3.2. Contribution of the 5HT1_A_ receptor

Administration of WAY (0.1, 1.0 mg/kg,) 15 minutes prior to the initiation of inhalation reliably attenuated the hypothermia induced by CBD [Time after vapor initiation, F (9, 54) = 54.31; P<0.0001; WAY dose, F (2, 12) = 8.185; P=0.0057; Interaction, F (18, 108) = 1.756; P=0.0405], as is shown in Figure 7A. The post-hoc test confirmed that temperature was significantly lower than in the Pre timepoint for the saline (30-240 minutes after start of inhalation), WAY 0.1 (30-240 minutes) and WAY 1.0 (60-240 minutes) pretreatment conditions. The post-hoc test further confirmed that body temperature was significantly different between saline and WAY 1.0 mg/kg pretreatment conditions from 30-240 minutes after the start of inhalation and between all three conditions from 60-120 minutes after the start of inhalation. Locomotor activity (Figure 7B) was significantly affected by Time after vapor initiation (F (9, 54) = 28.16; P<0.0001) and the interaction of Pretreatment condition with Time (F (18, 108) = 1.73; P<0.05). The post-hoc test confirmed that the 1.0 mg/kg WAY dose reduced locomotor activity in the Pre and 30-minute time-points relative to the other pre-treatment conditions.

**Figure 7:**
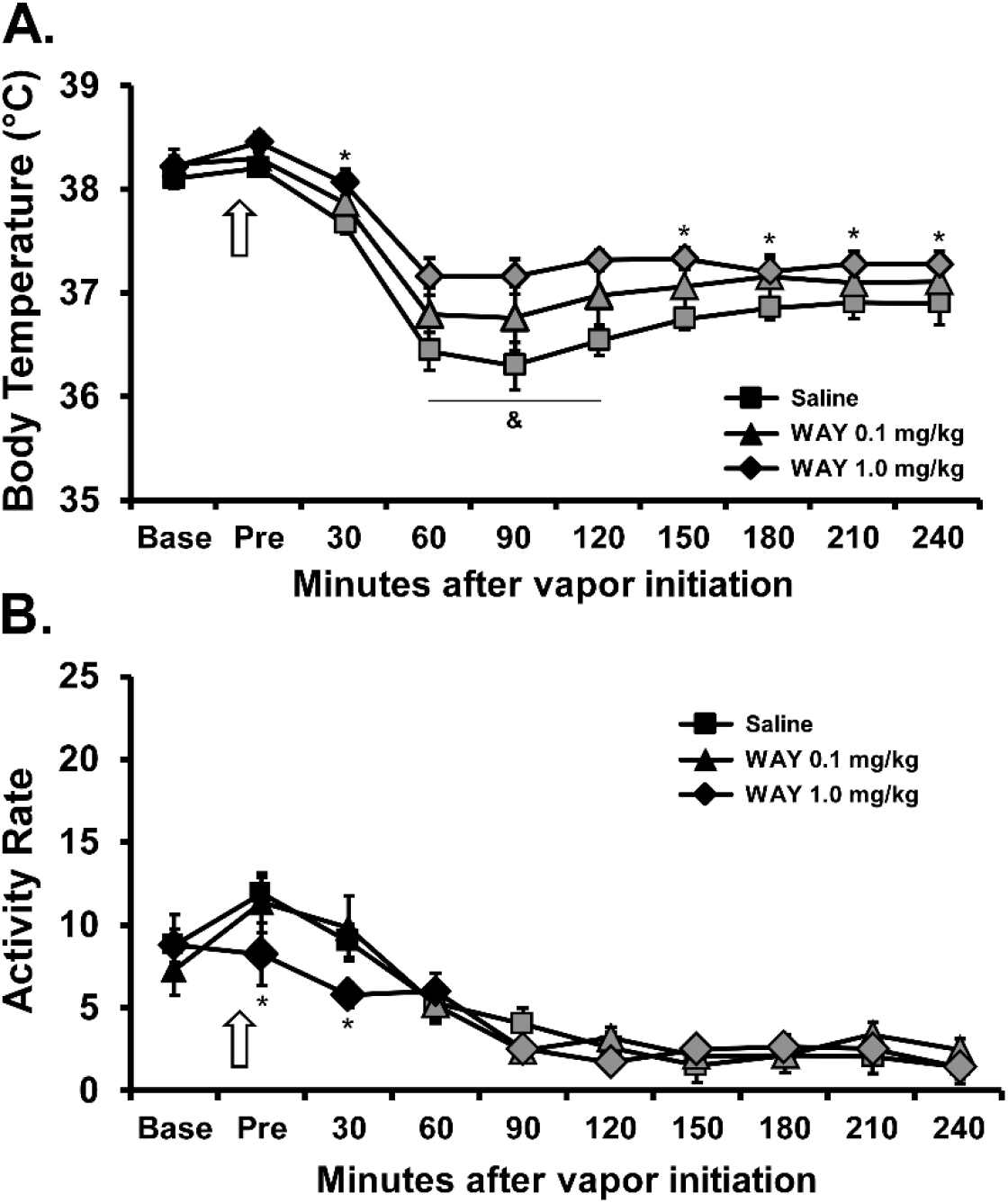
Mean (N=7; SEM) A) body temperature and B) activity rate following inhalation of CBD 100 mg/mL for 30 minutes after i.p. injection of WAY 100635 (0.1, 1.0 mg/kg) or the Saline vehicle. The arrow indicates the time of the pre-inhalation injection. Shaded symbols indicate a significant difference from the Pre-inhalation time-point, within treatment condition. Significant difference between all conditions is indicated with &, between the WAY 1.0 and the Saline pre-injection with *.

### 3.3 Experiment 4: CBD Effects on THC-induced Anti-nociception

In the first experiment, the inhalation of vapor from CBD (400 mg/mL) for 30 minutes (Figure 8A) had no impact on tail withdrawal latency, while the inhalation of THC 100 significantly slowed tail withdrawal [Pre/Post: F (1, 5) = 15.98; P<0.05; Vapor Condition: F (1, 5) = 5.39; P=0.068; Interaction: F (1, 5) = 7.79; P<0.05]. The threshold was confirmed at the 50 mg/mL concentration of THC [Pre/Post: F (1, 5) = 48.00; P<0.005; Vapor Condition: F (1, 5) = 2.64; P=0.1654; Interaction: F (1, 5) = 8.33; P<0.05] in the second experiment (Figure 8B). Finally, the analysis of the third experiment (Figure 8C) failed to confirm any significant alteration [Pre/Post: F (1, 5) = 33.09; P<0.005; Vapor Condition: F (1, 5) = 5.52; P=0.066; Interaction: F (1, 5) = 4.77; P=0.081] of the anti-nociceptive effect of THC (50 mg/mL) inhalation by co-inhalation with CBD (200 mg/mL).

**Figure 8:**
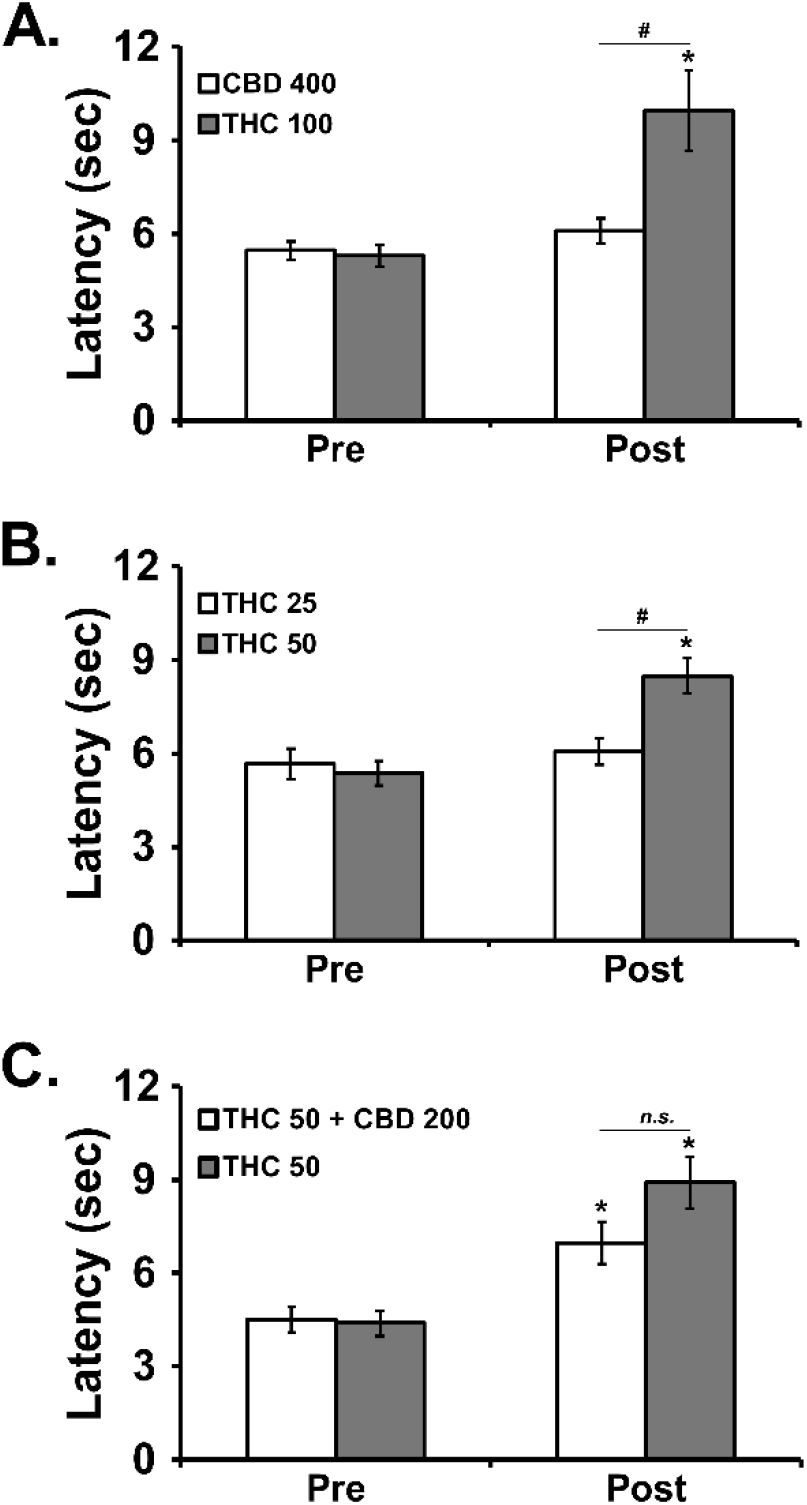
Mean (±SEM) tail withdrawal latency before (Pre) and after (Post) vapor inhalation for 30 minutes. Conditions included CBD (50, 400 mg/mL), THC (25, 50, 100 mg/mL) and the CBD (200 mg/mL)/THC (50 mg/mL) combination. A significant difference between Pre-vapor and Post-vapor is indicated with *, and a significant difference between inhalation conditions after vapor inhalation is indicated with #.

## 4. Discussion

The primary finding of this study was the confirmation that vapor inhalation using an e-cigarette based system elevates the plasma levels of CBD in male or female rats in a dose (i.e., concentration in the PG) dependent manner. Furthermore, it was shown that the CBD levels obtained in rats after 30 minutes of inhalation of vapor generated from concentrations of 100 to 400 mg/mL CBD in the PG vehicle fall within the range of the plasma levels produced 35 minutes after i.p. injection of 10 and 30 mg/kg CBD. The peak plasma concentrations were also similar to those reported for day old piglets after 1.0 mg/kg CBD, i.v. (Barata et al., 2019) but higher than those reported in humans (~62 ng/mL) using a plant material vaporizer with 11%THC/11%CBD cannabis (Arkell et al., 2019). The observed plasma concentrations were also comparable to plasma CBD observed in rats four hours after dermal gel application, which was the same time post-application that CBD reduced alcohol and cocaine seeking (Gonzalez-Cuevas et al., 2018). Only minor sex differences were observed, with slightly lower plasma CBD concentrations observed in the male rats after vapor inhalation or after a mg/kg adjusted i.p. injection. The larger body size of the males suggests perhaps a slightly different distribution per gram of bodyweight even when doses are adjusted for bodyweight when injecting. If anything, the differences in route of administration show a greater variability subsequent to i.p. injection versus inhalation (see error bars on Figure 2). Significantly reduced, but measurable, CBD was observed two hours after injection or the start of inhalation, echoing the elimination time-course observed in humans (Arkell et al., 2019). We have shown previously that 30 minutes of inhalation of vapor generated from CBD (100, 400 mg/mL) reduces the body temperature of male and female Wistar rats in a concentration-dependent manner (Javadi-Paydar et al., 2018a). This study replicated the thermoregulatory impact, thus the plasma levels obtained by vapor inhalation of CBD are those that produce robust physiological effects in the rat.

The radio-telemetry studies replicated our prior observation in Wistar male rats (Figure 3) and extended this by confirming that CBD inhalation reduces body temperature of male Sprague-Dawley rats (Figure 6) and that dose-dependent effects are obtained. We previously observed a greater sensitivity of Sprague-Dawley male rats compared with Wistar male rats to hypothermia induced by THC inhalation despite similar plasma THC levels and similar antinociceptive responses in a direct comparison study, available as a pre-print report (Taffe et al., 2019). A similar strain difference in the nadir of body temperature observed after inhalation of each concentration of CBD was observed in this study but the positive control experiment with the 5-HT_1A_ agonist 8-OH-DPAT, and the 5-HT_7_ antagonist SB-269970, confirmed that the Wistar rats were capable of a hypothermic response mediated by two key serotonergic contributors (Hedlund et al., 2004). This is consistent with the interpretation that Wistar male rats are less prone to reductions in body temperature, i.e., under threshold conditions that trigger a response in Sprague-Dawley male rats. This may be a result of their larger body size, relative fat/lean body mass distribution or some other factor.

The 5-HT_1A_ receptor antagonist (WAY-100,635; WAY) had a reliable and dose-dependent, but limited, effect on the hypothermia produced by CBD inhalation with effects observed only in the Sprague-Dawley group. The failure to observe any effect in the Wistar group was not due to a lack of contribution of 5-HT_1A_ to thermoregulation in this strain since injection of the agonist produced a profound hypothermia that was opposed by the same doses of WAY that were inactive against CBD inhalation. The blunted response of the Wistar rats to CBD (100 mg/mL) inhalation in the antagonist study, compared with their first response to the same dose further complicates interpretation since they had possibly developed some degree of tolerance. Since the magnitude of the WAY attenuation in the Sprague-Dawley rats was modest (Figure 7), strong conclusions about a differential role of 5-HT_1A_ in CBD vapor-induced hypothermia across strains is premature. Relatedly, the suppression of locomotor activity associated with WAY 1.0 mg/kg pre-treatment the Sprague-Dawley group confirmed that an active dose was selected and this enhances confidence that the result was not merely a question of an inactive WAY dose. Overall, these data confirm that the thermoregulatory response to CBD vapor inhalation is mediated by the 5-HT_1A_ receptor subtype, but only partially so. Additional investigation would be required to determine the additional pharmacological contributors; obviously 5-HT_7_, TRPV2 and CB_1_ receptors would be potential targets for further study, as reviewed (Boggs et al., 2018).

Interestingly, the Sprague-Dawley male group was tolerant to the hypothermic effects of nicotine inhaled alone at the 30 mg/mL concentration, compared with their initial responses in a previously published study (Javadi-Paydar et al., 2019), but nevertheless exhibited a nicotine-associated enhancement of the hypothermia caused by CBD at either vapor concentration. Locomotor effects of nicotine inhalation were observed in this study and showed that an active dose was administered, despite evidence of tolerance to the hypothermic effects. Interestingly, CBD was able to suppress the locomotor stimulant effect of nicotine when the two were administered in combination. CBD inhaled by itself did not reduce locomotor activity below that observed after PG inhalation in the Sprague-Dawley rats (Figure 6C,D), unlike the small locomotor suppression that was observed after CBD inhalation in both male and female Wistar rats in a prior report (Javadi-Paydar et al., 2018a) and in the male Wistar rats in this study (Figure 3B). Locomotor responses are not a precise translational match for abuse liability, or treatment thereof, but these results at least align with a preliminary double-blind placebo controlled study found that CBD delivered by inhaler was able to reduce cigarette consumption over a week of treatment (Morgan et al., 2013). As with our prior study of the effects of nicotine and THC co-inhalation (Javadi-Paydar et al., 2019), the effects on locomotion and body temperature are most consistent with independent effects of CBD and nicotine, which are in the same direction for body temperature but in opposition for locomotor stimulation.

In the nociception experiment, THC increased tail-withdrawal latency after inhalation of vapor for 30 minutes in the 50-100 mg/mL concentrations but not at the 25 mg/mL concentration. CBD did not significantly alter nociception by itself or significantly alter the anti-nociceptive effect of THC (50 mg/mL). These findings align with prior research examining the parenteral injection of CBD. For example, CBD is inactive in a tail withdrawal assay in rat when injected by itself or when co-injected with THC, despite enhancing the effects of THC on paw pressure nociception (Britch et al., 2017). Cannabidiol, i.c.v., does enhance the anti-nociceptive effects of i.c.v. morphine using the warm water tail-withdrawal method in mice (Rodriguez-Munoz et al., 2018), but CBD is not effective in the tail-withdrawal assay in mice when injected i.p. alone or in combination with morphine (Neelakantan et al., 2015). CBD reduced carrageenan-induced hyperalgesia (in a CB_1_ and CB_2_ dependent manner) and acetic acid-induced writhing (in a CB_1_ independent manner), but not hot-plate analgesia, in mice (Silva et al., 2017). There is a clear need for broader study of the effects of CBD, including across routes of administration. CBD inhalation reduces body temperature in rats, in a manner that depends in part on the 5-HT_1A_ receptor subtype, but CBD does not appear to do so when administered by parenteral injection (Taffe et al., 2015). This might suggest that in some cases the route of administration is crucial to the impact of CBD, making this a major translational consideration since inhalation, transdermal and oral routes are already common in human user populations.

In conclusion, these data confirm a method for studying the effects of inhaled CBD in a rat model and, together with our prior reports (Javadi-Paydar et al., 2018a; Taffe et al., 2015), suggest that the route of administration may confer different effects *in vivo* even with approximately similar plasma CBD exposure.

## Acknowledgements

These studies were supported by USPHS grants R01 DA035281; R01 DA035482; R44 DA041967 and R01 DA042595. The NIH/NIDA had no role in study design, collection, analysis and interpretation of data, in the writing of the report, or in the decision to submit the paper for publication. The authors are grateful to Sophia A. Vandewater for expert technical assistance with several of the experiments. La Jolla Alcohol Research, Inc (LJARI) engages in commercial development of vapor inhalation techniques and equipment, including with support from the R44 DA041967 SBIR grant. LJARI was not directly involved in the design of the experiments, analysis and interpretation of data or the decision to submit the study for publication. The authors declare no additional financial conflicts which affected the conduct of this work.

## Notes

#### Summary of Updates

This revised version includes a new study in Wistar male rats.

